# Compartmentalized RNA catalysis in membrane - free coacervate protocells

**DOI:** 10.1101/273417

**Authors:** Björn Drobot, Juan M. Iglesias-Artola, Kris Le Vay, Viktoria Mayr, Mrityunjoy Kar, Moritz Kreysing, Hannes Mutschler, T-Y. Dora Tang

## Abstract

Phase separation of mixtures of oppositely charged polymers provides a simple and direct route to compartmentalization via coacervation, which may have been important for driving primitive reactions as part of the RNA world hypothesis. However, to date, RNA catalysis has not been reconciled with coacervation. Here we demonstrate that RNA catalysis is viable within coacervate microdroplets and further show that these membrane-free droplets can selectively retain longer length RNAs while permitting transfer of lower molecular weight oligonucleotides.

## Introduction

Compartmentalization driven by spontaneous self-assembly processes is crucial for spatial localization and concentration of reactants in modern biology and may have been important during the origin of life. One route known as coacervation describes a complexation process^1,2^ between two oppositely charged polymers such as polypeptides and nucleotides.^3–7^ The resulting coacervate microdroplets are membrane free, chemically enriched and in dynamic equilibrium with a polymer poor phase. In addition to being stable over a broad range of physicochemical conditions, coacervate droplets are able to spatially localize and up-concentrate different molecules^3,8^ and support biochemical reactions.^9,10^

It has been hypothesized that compartments which form via coacervation could have played a crucial role during the origin of life by kick-starting and concentrating the first biochemical reactions on Earth.^11^ Coacervation has also been implicated in modern biology where it has been shown that the formation of membrane-free compartments or condensates such as P-bodies or stress granules within cells are driven by this mechanism.^12,13^ These membrane free organelles are chemically isolated from their surrounding cytoplasm through a diffusive phase boundary, permitting the exchange of molecules with their surroundings.^14^ In addition, these compartments may localize specific biological reactions and play important roles in cellular functions such as spatial and temporal RNA localization within the cell.^15–18^

Whilst there is increasing evidence for the functional importance of RNA compartmentalization via coacervation in modern biology, this phenomenon would also have been vitally important during a more primitive biology. Up-concentration and localization could have enabled RNA to function both as a catalyst (ribozyme) and storage medium for genetic information, as required by the RNA world hypothesis.^19^ To date, ribozymes have been encapsulated within eutectic ice phases^20,21^ and protocell models such as water-oil-droplets for directed evolution experiments,^22–24^ membrane-bound lipid vesicles,^25–27^ and membrane free compartments based on PEG/Dextran aqueous two-phase systems (ATPS).^28^ Interestingly, RNA catalysis within ATPS exhibits an increased rate of reaction as a result of the increased concentration within the dextran phase. Despite these examples, RNA catalysis has not been demonstrated within coacervate based protocells. Herein, we show the ability of the coacervate microenvironment to support RNA catalysis whilst selectively sequestering ribozymes and permitting transfer of lower molecular weight oligonucleotides.

## Results & Discussion

We developed a real-time fluorescence resonance energy transfer (FRET) assay (see materials and methods and ESI) to investigate the effect of the coacervate microenvironment on catalysis of a minimal version of the hammerhead ribozyme derived from satellite RNA of tobacco ringspot virus (HH-min)^29^. HH-min and its FRET-substrate (Figure 1A, materials and methods and ESI) were incubated within a bulk polysaccharide / polypeptide coacervate phase or within coacervate microdroplets (see ESI) under single turnover conditions. Cleavage of the FRET-substrate strand by HH-min increases the distance between 6-carboxyfluorescein (FAM) and Black Hole quencher 1 (BHQ1), resulting in increased fluorescence intensity. We further developed an inactive control ribozyme (HH-mut) by introducing two point mutations at the catalytic site (see materials and methods and ESI).

**Figure 1.**
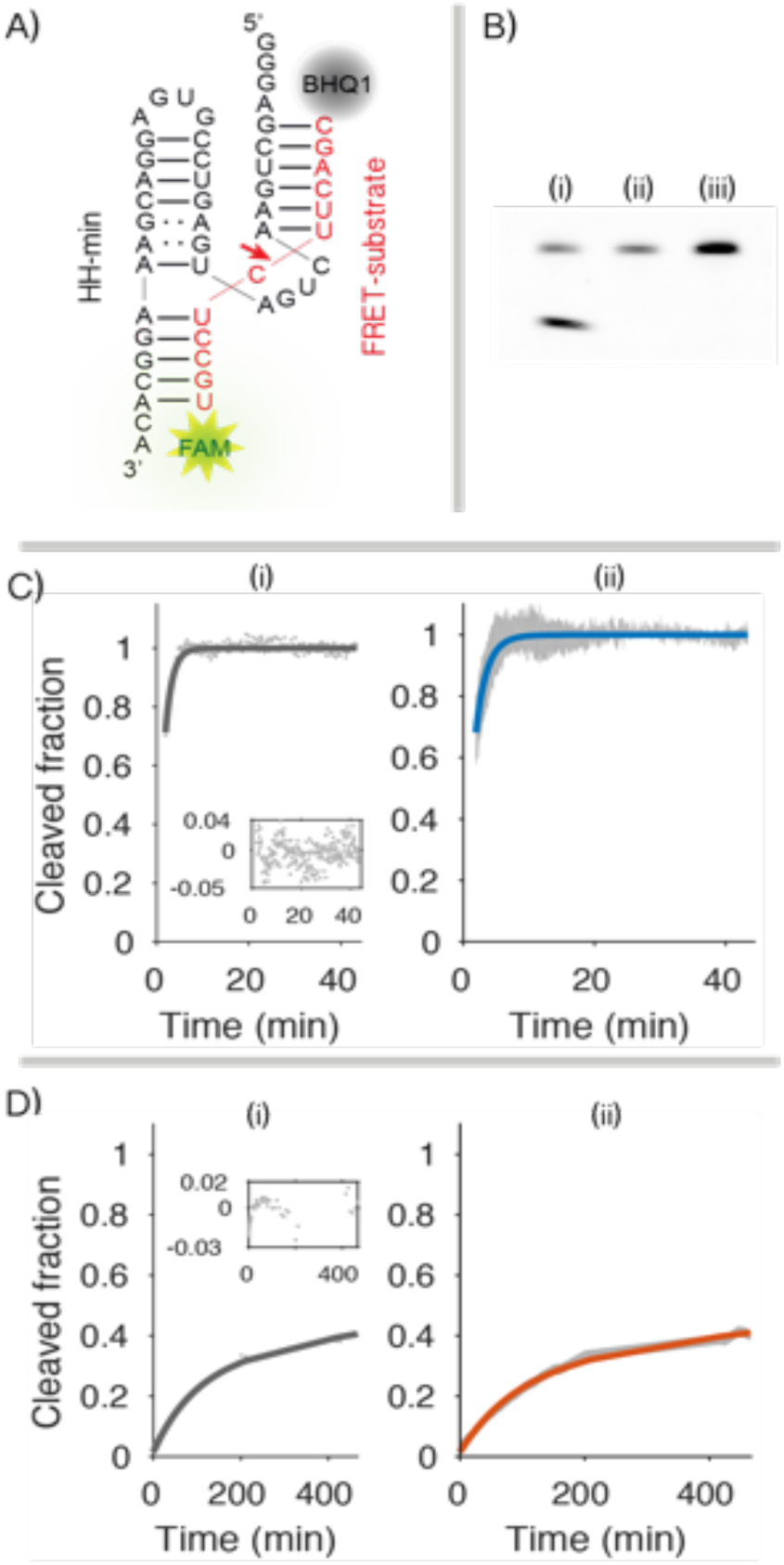
Cleavage of the FRET-substrate under different conditions. **(A)** HH-min (black) and the FRET substrate (red). **(B)** Gel electrophoresis of RNA cleavage in bulk coacervate phase (4:1 CM-Dex:PLys). 0.5 µM of FRET-substrate was incubated with 1 µM of (i) HH-min, (ii) HH-mut or (iii) no ribozyme in bulk phase (25 °C, 60 min). Samples were analyzed by denaturing PAGE followed by fluorescence imaging. The lack of in-gel quenching of the FRET substrate likely results from modifications of BHQ1 during PAGE.^41^ **(C)** Real-time cleavage kinetics in 10 mM Tris•HCl pH 8.3 and 4 mM MgCl_2_. (i) A monoexponential fit (grey line) to kinetic data (grey dots) and residuals of the fit (inset) (ii) Blue line is the mean of the individual fits (blue line). Grey data points represents the standard deviation (N ≥ 5) from experimental data. **(D)** Cleavage in bulk coacervate phase (normalized to the amount of cleaved product at t = 530 min from gel electrophoresis). (i) Biexponential fit (dark grey line) to experimental data (grey dots) with the residuals (inset) (ii) mean biexponential fit (orange) of individual fits (N ≥ 5). Grey data points represents the standard deviation (N ≥ 5) from the experimental data.

HH-min (1 µM) and FRET-substrate (0.5 µM) were incubated within the CM-Dex:PLys bulk coacervate phase (4:1 final molar concentration, pH 8.0). The RNA was then separated from the coacervate phase and analyzed by denaturing gel. Excitingly, fluorescence gel imaging showed the presence of cleavage product in the bulk coacervate phase containing HH-min. In contrast, control experiments in the absence of HH-min or in the presence of HH-mut showed no evidence of the cleavage product, confirming that the wild type ribozyme drives substrate cleavage in the bulk coacervate phase (Figure 1B). The FRET assay was further exploited to characterize the enzyme kinetics in both the bulk coacervate phase and buffer by time resolved fluorescence spectroscopy under single turnover conditions by direct loading of HH-min and FRET-substrate into either cleavage buffer or bulk coacervate phase. The increase in fluorescence intensity of FAM was measured over time and normalized to the amount of cleaved product produced (see ESI). Fitting the kinetic profiles of substrate cleavage in buffer conditions with a single exponential revealed an apparent rate constant, k_0_ of 0.6 +/-0.2 min^-1^ (Figure 1C) which was comparable to the k_0_ obtained in buffer analyzed by gel electrophoresis (0.4 ± 0.01 min^-1^) (Figure S1) and to k_cat_ values previously determined for a range of hammerhead ribozymes (0.01-1.5 min^-1^).^29,30^ In comparison, RNA cleavage within the bulk coacervate phase was clearly biphasic (Figure S2A) with an observed faster rate constant of k_1_ of 1.0 × 10^−2^ +/-0.1 × 10^−2^ min^-1^ and a second slower rate constant of k_2_ of 4.0 × 10^−4^ +/-1.0 × 10^−4^ min^-1^. Thus, the fastest rate constant k_1_ is an order of magnitude slower than in buffer conditions (k_0_ = 0.6 +/-0.2 min^-1^) indicative of reduced activity within the coacervate phase. In addition, the transition to biphasic kinetics within the coacervate phase compared to the aqueous buffer phase describes a different kinetic mechanism of HH-min within the coacervate phase (Figure 1D). This may be attributable to heterogeneous ribozyme populations with alternative conformational and equilibrium states, as observed for some HH systems in aqueous buffer conditions.^31,32^ It is possible that the charged and crowded coacervate microenvironment affects the structure of HH-min, restricting substrate binding, sterically hindering substrate-enzyme complex formation and/or spatially restricting diffusion of the cleavage assay components. Indeed, measured diffusion coefficients of TAM-HH-min (1.0 +/-0.2 µm^2^·s^-1^) and FAM-substrate (1.6 +/-0.1 µm^2^·s^-1^) in the bulk coacervate phase (Figure 2) phase from FRAP analysis showed a significantly slower molecular diffusion of the ribozyme and substrate compared to predicted diffusion coefficients of RNA in buffer (∼150 µm^2^·s^-1^, Figure 2).^33,34^ The decreased mobility is indicative of a more viscous and spatially restricted environment in the interior of the coacervate phase (η = 114 +/-21 mPa·s, Figure 2C).

**Figure 2.**
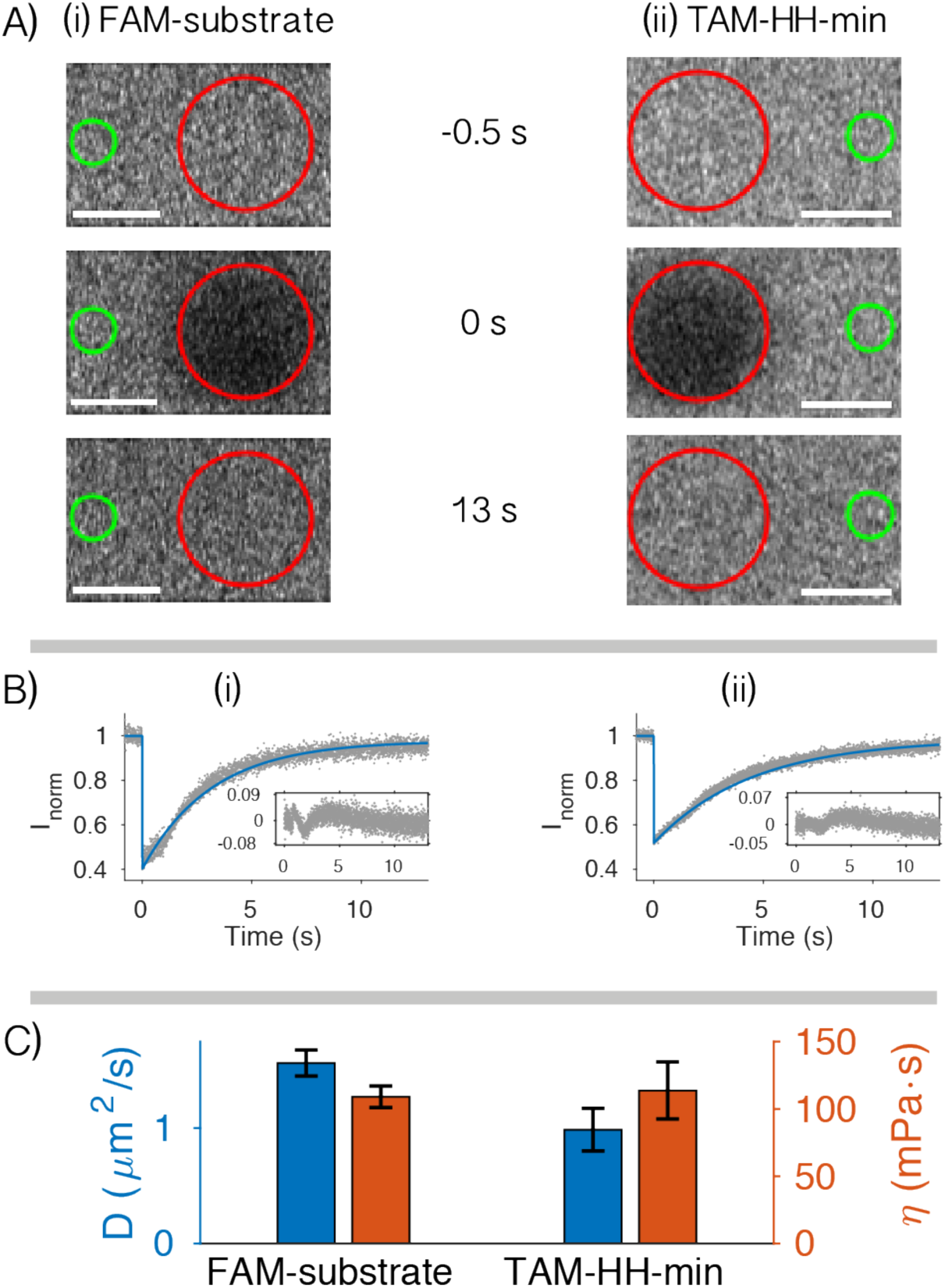
FRAP of bulk coacervate phase. Bulk phase (4:1 CM-Dex:PLys) containing either (i) 0.36 µM FAM-substrate or (ii) 0.36 µM TAM-HH-min (ii). **(A)** Output frames from confocal imaging (63x) are shown at t = −0.5 s before bleaching, directly after bleaching (red circle, t = 0 s) and t = 13 s after bleaching. The fluorescence intensity was normalized against a reference (green circle) and fit to standard equations. Scale bars are 5 µm. **(B)** Plots of normalized FRAP data for HH-min (ii) and FAM-substrate (ii) show the standard deviation (grey, N=10) and fit (blue) from the same bleach spot radius. **(C)** Diffusion coefficients and viscosities obtained from (B). Mean and standard deviations are from at least two different samples with analysis from ≥ 14 bleach spots for each experiment.

To test the activity of the ribozyme within individual droplets, the bulk coacervate phase containing ribozyme and substrate was re-dispersed in supernatant to produce microdroplets in solution (see materials and methods). The final concentration of enzyme and substrate in the microdroplet dispersion was equivalent to the final concentration of the bulk coacervate phase under single turnover conditions (1 µM of HH-min and 0.5 µM FRET-substrate). Fluorescence optical microscopy images showed an increase in FAM fluorescence intensity in the droplets after 900 min (Figure 3A).

**Figure 3.**
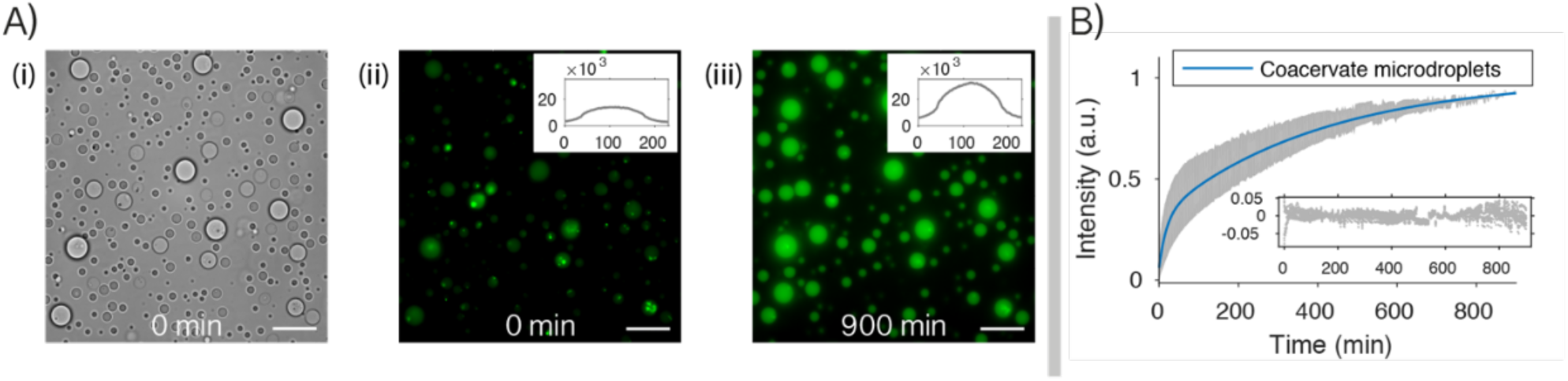
RNA catalysis in coacervate microdroplets. **(A)** (i) Optical microscopy images of 4:1 CM-Dex:PLys coacervates prepared in cleavage buffer (1 µM of HH-min and 0.5 µM FRET-substrate). Fluorescence microscopy images at t = 0 min (ii) and t = 900 min show an increase in FAM fluorescence (see inset). Scale bars are 20 µm. **(B)** Background corrected and volume / endpoint normalized fluorescence intensity of droplets. Standard deviation of kinetics from twelve micro-droplets (grey) with the mean biexponential fit (blue) and residuals (inset).

Fitting the biphasic fluorescence signal allowed a direct comparison of kinetic profiles between the coacervate microdroplet (Figure 3B, Figure S2B) and bulk coacervate phase environments. A modestly faster rate constant (k_1_ and k_2_) was observed in the microdroplets (k_1_ of 4.8 × 10^−2^ +/-0.1 × 10^−2^ min^-1^, k_2_ of 2.4 × 10^−3^ +/-0.2 × 10^−3^ min^-1^) compared to the bulk coacervate phase (k_1_ of 1.1 × 10^−2^ +/-0.2 × 10^−2^ min^-1^, k_2_ of 4.0 × 10^−4^ +/-1.0 × 10^−4^ min^-1^) (Table S3, Figure S3). Quantitative FRAP analysis, showed that the measured viscosities of these two environments are the same within error (Figure S4). Given these conditions, it would be expected that k_1_ and k_2_ would be the same in both the coacervate bulk phase and within the coacervate microdroplets, however a slight increase in the rate constant was observed. Determination of the partition coefficients of both the ribozyme and substrate (K_HH-min_ = 7150 +/-1650 and K_HH-substrate_ = 1280 +/-200) (Figure S5) by fluorescence spectroscopy (see ESI) showed that both TAM-HH-min and FAM-substrate partition strongly into the coacervate environment. This could lead to the possibility of a slight alteration of the material properties within the coacervate microdroplets providing a straightforward explanation to the observed variations in the rate constants.

To further investigate the six-fold difference in the partition coefficient between the ribozyme and substrate, we characterized the differences in the rate and extent of sequestration of TAM-HH-min and FAM-substrate from the surrounding aqueous phase into the droplet after whole-droplet photobleaching. Coacervates containing FAM-substrate showed complete fluorescence recovery within 100 s and a recovery half time (t) of 22 +/-0.1 s. In comparison, TAM-HH-min showed only 70 % recovery after 300 s with t = 189 +/-14 s, attributed to either a low concentration of HH-min within the surrounding aqueous phase and/or a slow rate of transfer into the coacervate droplet from its exterior (Figure S6). Taken together, these results show that the 12-mer substrate has a lower affinity for the coacervate droplet and a faster exchange between the droplet and the surrounding aqueous phase compared to the 26-mer ribozyme.

Our results, and others,^4^ show that RNA sequestration and localization within coacervate droplets is dependent on the length of the RNA. Selective retention of longer length RNA with transfer of shorter length RNA can thus have implications for ribozyme catalysis within coacervate droplets. To investigate this, we directly observed the localization of TAM-HH-min and FRET-substrate. CM-Dex:PLys coacervate micro-droplets (4:1 final molar ratio) containing TAM-HH-min were loaded into one end of a capillary channel (Figure 4A, region 1) while droplets containing the FRET-substrate were loaded into the other end of the channel (Figure 4A, region 4). Time resolved fluorescence optical microscopy images in both the TAM and FAM channels were obtained at various locations along the capillary channel (Figure 4A, regions 1, 2, 3). Imaging over 500 min in the TAM channel showed that, within the measurable resolution, no diffusion of the ribozyme to droplets in other regions of the channel occurs (Figure 4B (2) and S8). Conversely, over the time course droplets in all three regions exhibited increased FAM fluorescence with droplets in region 1 with the highest intensity and droplets in region 2 and 3 with comparatively lower intensity (Figure 4C). Analysis of time resolved images showed a delayed increase in the onset of cleaved product fluorescence in region 2 and a further delayed onset in cleaved product in region 3. These results are commensurate with diffusion of the FRET-substrate out of the droplets in region 3 and into droplets containing TAM-HH-min in region 1 where cleavage takes place. The cleaved product then diffuses out of the active droplets and into droplets in regions 2 and 3. Taken together, our results show that longer length RNA (HH-min) is retained and spatially localized within the highly charged and crowded interior of the coacervate droplet, while shorter RNAs transfer between droplets.

**Figure 4.**
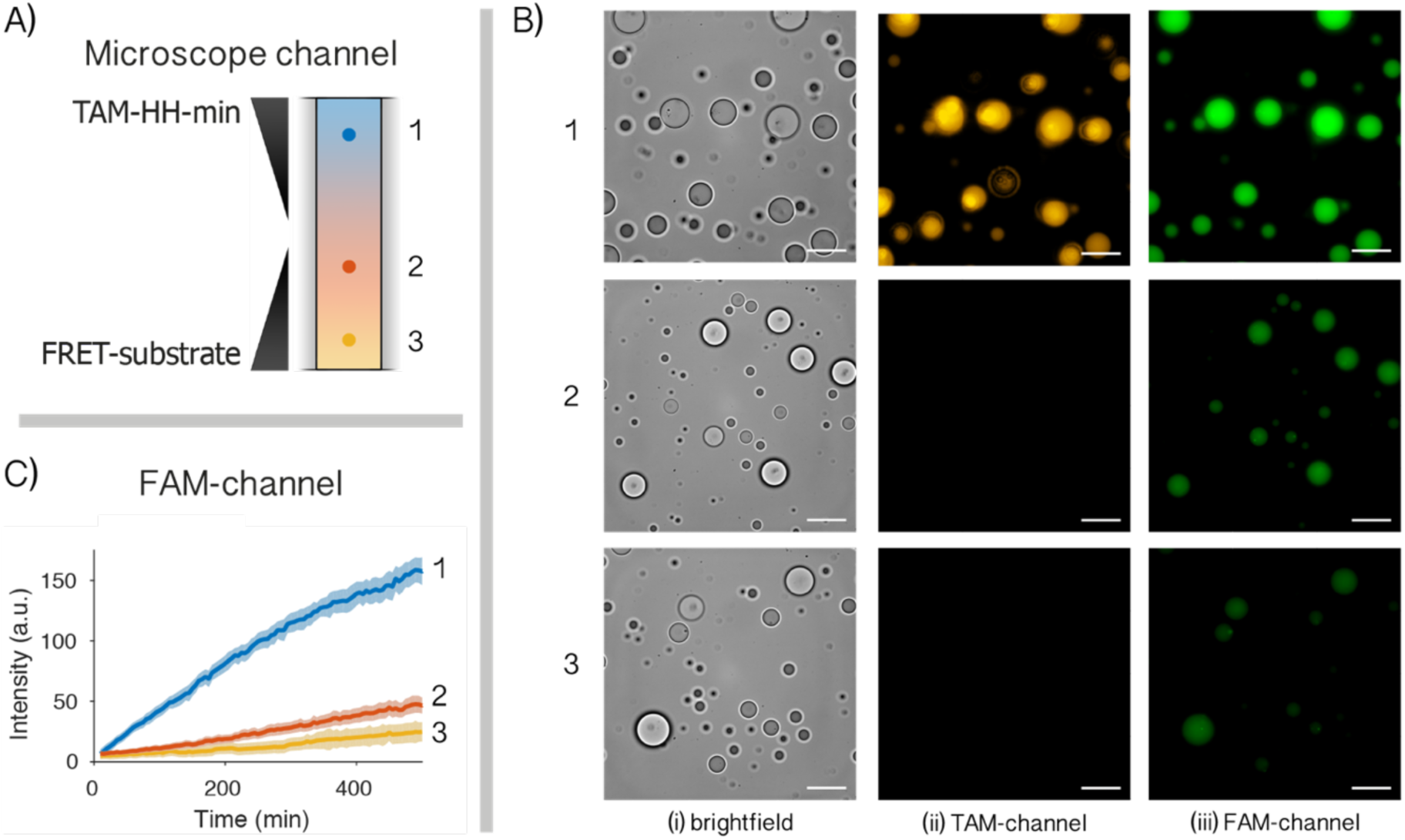
Localization and retention of RNA within coacervate droplets. **(A)** Schematic of localization experiments where 4:1 CM-Dex:PLys coacervate droplets containing 0.36 µM (final concentration) TAM-HH-min were loaded into one end of a capillary channel (1). Droplets containing 0.36 µM FRET-substrate were loaded into the other end of the channel (3). **(B)** Optical microscopy images obtained using a 100x oil immersion lens in (i) bright field and fluorescence mode using filters for (i) TAM or (ii) FAM (iii). Images were captured in regions 1, 2 and 3 at t = 500 min (scale bar: 20 µm). **(C)** FAM fluorescence intensity kinetics of the cleaved substrate in at least seven droplets from region 1 (blue), 2 (orange) and 3 (yellow).

## Conclusions

Here, we demonstrate that coacervate microdroplets offer intriguing properties for compartmentalized RNA catalysis. Our results show that these membrane-free microenvironments support RNA catalysis and up-concentrate oligonucleotides within their interiors. Moreover, this is coupled to selective retention and release of RNA without additional energy input. These features could have been significant on early Earth where concentrations of RNA and their building blocks may have been low. Moreover, maintenance of the genetic identity of coacervate protocells could be achieved via spatial localization of RNA catalysis and RNA genomes with spread of RNA building blocks or short genetic polymers between droplets. Whilst this work represents a key step in reconciling primitive RNA catalysis with selective protocellular compartmentalization it should also be noted that these features of compartmentalization are significant in modern biology. To this end, there are immediate questions to be addressed. For example, how does the microdroplet environment alter the ribozyme mechanistic pathway and effect nucleotide selectivity into the droplet? How can we alter the physical chemistry of the droplets to affect oligonucleotide selectivity? In addition, our experiments have focused on a nucleolytic ribozyme, however rather than RNA cleavage an increase in genetic and molecular complexity of RNA e.g. by ligation would have been important during early Earth. Therefore, probing RNA synthesis through ligase activity within coacervate microdroplets will further contribute to the understanding of the role of coacervation on early Earth and modern biology.

## Materials & Methods

### RNA synthesis

A minimal, *trans*-acting hammerhead ribozyme (HH-min) derived from satellite RNA of tobacco ringspot virus and complementary substrate were produced by modification of the helix 1 hybridizing arm in a *cis-*acting system.^35^ An inactive variant (HH-mut) was produced by the introduction of two point mutations, G5A and G12A, which inhibit correct ribozyme folding and active site protonation-deprotonation events respectively.^36^ The wild-type and inactive hammerhead ribozymes were transcribed *in vitro* by T7 RNA polymerase. The DNA templates for transcription were produced by fill-in of DNA oligonucleotides using GoTaq (Promega). Complementary pairs of DNA oligonucleotides contained the ribozyme gene (sTRSV_min_wt_TX/ sTRSV_min_mut_TX) and T7 promoter (5T7) (Table S1). The double-stranded templates were purified using a Monarch PCR DNA Cleanup Kit (NEB, Biolabs, USA). Ribozyme RNA was transcribed from the double-stranded templates using the MEGAshortscript(tm) T7 Transcription Kit (ThermoFisher), and purified using RNeasy (Qiagen).

### Preparation of bulk coacervate phase and coacervate microdroplets containing RNA HH and substrate

To prepare bulk coacervate phase containing ribozyme and substrate, aqueous dispersions of 4:1 molar ratio of CM-Dex: PLys coacervate microdroplets in 10 mM Tris•HCl pH 8.0 and 4 mM MgCl_2_ were prepared by mixing 250 µl of 1 M CM-Dex, 200 µl of 0.2 M PLys, 50 µl of 1 M Tris•HCl pH 8.0, 20 µl of 1 M MgCl_2_ and made up to 5 ml with nuclease free water. The bulk polymer phase was produced by separating the droplets from the supernatant phase by centrifugation (10 min, 4000 g) and removing the supernatant. Coacervate microdroplets containing ribozyme or substrate were prepared by mixing RNA with 1 µl of the coacervate phase and then redispersing the polymer phase in 49 μl supernatant by vortexing.

### Hammerhead activity within bulk coacervate phase or coacervate microdroplets by Gel electrophoresis

HH-min and FRET-substrate at final concentrations of 1 µM and 0.5 µM respectively were loaded into bulk coacervate phase for single turnover conditions. RNA was extracted from 5 µl of bulk coacervate phase (25 °C) or from 5 µl of coacervate microdroplet dispersions after 900 min (25 ^o^C) by sequential addition, 1 s of vortexing and 1 s of centrifugation of 5 µl of 1 M NaCl (final concentration 4.8 mM), 5 µl of 1.25 M Hexametaphosphate (final concentration - 6.0 mM) and 90 µl RNA loading buffer (final volume - 83 %) containing EDTA (final concentration - 8 mM) and Orange G. The reaction mixture was heated at 80 °C for 10 min, centrifuged and placed on ice for at least 5 min. 10 µl of the reaction mixture was loaded into a pre-run 20% polyacrylamide gel and then run at 300 V in 1x TBE buffer until the dye had run to the bottom of the gel. The gel was imaged using Typhoon FLA-9500, GE Healthcare Life Sciences with λ_ex_ = 473 nm, λ_em_= 520 nm. Band intensities were measured using FIJI to obtain the ratio of cleaved product at a specific time point, which was then used to normalize kinetic data obtained from spectroscopy or microscopy.

### Kinetics of ribozyme activity within bulk coacervate phase or buffer

The enzyme reaction was incorporated into either buffer or bulk coacervate phase by adding the appropriate volume from ribozyme stock (HH-min or HH-mut) solutions into the polymer phase to achieve a final concentration of 1 µM. The reaction was initiated by adding FRET-substrate at a final concentration of 0.5 µM. After mixing the reaction mixture, 20 µl of samples were loaded into a 384 well plate (microplate, PS, Small Volume, LoBase, Med. binding, Black, Greiner Bio-one). The enzyme activity was monitored using TECAN Spark 20 M well plate reader spectrophotometer (Tecan AG, Männedorf, Switzerland) by measuring the increase in FAM fluorescence over time (λ_exc_ = 485 nm and λ_em_ = 535 nm, 10 nm bandwidth, at 25 °C). The fluorescence intensity of HH-mut used as the background intensity and subtracted from HH-min data. This was then normalized by determining the amount of FRET-substrate cleaved by gel electrophoresis at the endpoint of the experiments as described previously. Kinetic profiles were fit to either single exponential growth (equation 1) under buffer conditions or bi-exponential growth (equation 2) for bulk coacervate experiments.

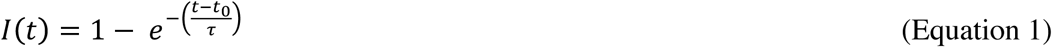

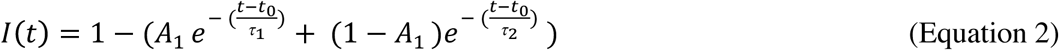

The corresponding rate constants k_1_ or k_2_ are obtained from the fitted time constants t_1_ or t _2_ where k=1/ t.

### Hammerhead catalysis within coacervate microdroplets

FRET based HH-min activity within coacervate droplets was undertaken by addition of the FRET-substrate into a dispersion of coacervate droplets containing ribozyme (final concentration under single turnover conditions were 1 µM for HH-min and 0.5 µM FRET-substrate and then loaded into custom made PEGylated channel capillary slide for microscope imaging. Time resolved bright field and fluorescent images of the droplets were obtained using a 100 × oil immersion objective (Plan-Apochromat 100x/1.40 Oil DIC, Zeiss) mounted onto a Zeiss Axiovert 200M inverted widefield microscope equipped with a 16-channel CooLED and an ANDOR ZYLA fast sCMOS camera. Image acquisition was controlled with the Metamorph software (1 frame / minute, 100 ms exposure time) for 900 min. Images were taken with λ_exc_ = 475 +/-28 nm and λ_em_ = 525 +/-15nm and analyzed using the Fiji software to obtain the integrated fluorescence intensities divided by the volume of the droplet as a function of time for 12 coacervate microdroplets. The amount of substrate cleaved was normalized by gel electrophoresis at a given time point as described previously. Kinetic parameters were derived from fitting the kinetic data to Equation 2. To test for reproducibility the experiment was repeated with a different batch of HH-min using the same experimental conditions.

### Determining the diffusion coefficient and the localization of RNA using Fluorescence Recovery after Photobleaching (FRAP)

Fluorescence recovery after photobleaching was undertaken within both coacervate microdroplets and the bulk coacervate phase containing either TAM-HH-min (0.36 µM) or FAM-substrate (0.36 µM). Samples were prepared as previously described and loaded into capillary slides mounted in a Zeiss LSM 880 inverted single point scanning confocal microscope equipped with a 32 GaAsP PMT channel spectral detector and a 32-channel Airy Scan detector and imaged using a 63x oil immersion objective (Plan-Apochromat 63x 1.4 Oil DIC, Zeiss). Bleaching was achieved by additional excitation with a 405 nm laser diode and 355 nm DPSS laser, an Argon Multiline Laser produced the excitation wavelength of λ_FAM_ = 488 nm or ? _TAM_ = 561 nm and emission wavelengths λ_FAM_ = 479 to 665 (beam splitter 488 nm) nm or λ_TAM_ = 562 to 722 nm. Imaging time varied depending on the region of interest but was typically between 12 ms/frame and 100 ms/frame.

The fluorescence intensity as a function of time for the bleached area, reference and the background were obtained using FIJI and the recovery of the bleached region was normalized against the background and the reference region for bulk coacervate experiments. An additional normalization for droplet based FRAP was undertaken by dividing the fluorescence by the fluorescence of the whole droplet^37,38^. The kinetic profiles were fit to Equation 3 using MATLAB to obtain the time constant, *t*, of fluorescence recovery or of transport into the droplet (whole droplet FRAP).

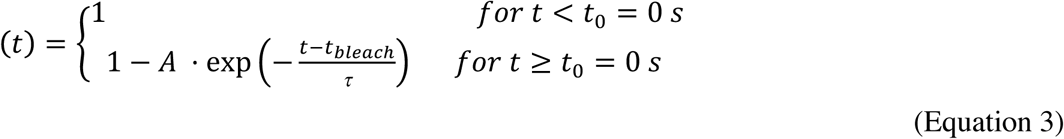

Where *t*_*0*_ = 0s is defined as the first time point after bleaching. The diffusion coefficient is related to the time constant *t* by the relation.^39,40^ The diffusion coefficient was averaged over twenty bleaching events across at least two different samples:

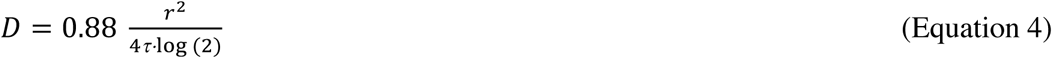

Here *r* is the radius of the bleached spot. The diffusion coefficient was averaged over twenty bleaching events across at least two different samples. Interpretation of the apparent diffusivity from photobleaching experiments may be complicated due to the complexity of the liquid coacervate phase including interactions between the studied fluorescence labelled RNA and the polymers. However, by approximating the coacervate phase as a Newtonian fluid an estimation of the viscosity via the Stokes-Einstein relation (Equation 5) could be made

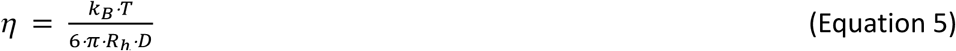

with *η* the viscosity (mPa·s), k_B_ the Boltzmann constant (m^2^kg s^-2^K^-1^), *T* the temperature in K and *R*_*h*_ the hydrodynamic radius (in m) calculated from length of the RNA using Equation 6^7^.

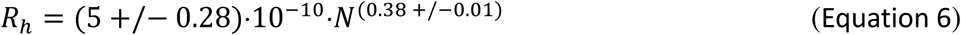

(Equation 6) where *N* is the length in nucleotides.

### Ribozyme activity and localization within coacervate droplets

To determine the localization of the RNA, 4:1 CM-DEX: PLys coacervate micro-droplets emulsions (1 µl bulk coacervate phase mixed with 49 µl supernatant) containing TAM-HH-min (0.36 µM) were loaded into one end of a capillary channel whilst droplets containing the FRET-substrate (0.36 µM) were gently loaded into the other end of the channel in such a way to prevent droplet mixing whilst permitting passive diffusion of molecules through the length of the channel. Control experiments also included loading droplets containing either TAM-HH-min or FAM-substrate into one end of the channel whilst microdroplets containing HH-DUB were loaded into the other end of the capillary channel.

Time resolved bright field and fluorescent images of the droplets along different parts of the imaging channel were obtained using a 100x oil immersion objective (Plan-Apochromat 100x/1.40 Oil DIC, Zeiss) mounted onto a Zeiss Axiovert 200M inverted widefield microscope equipped with a 16-channel CooLED and an ANDOR ZYLA fast sCMOS camera. Images in the TAM channel were taken with λ_exc_ = 542 +/-13.5 nm and λ_em_ = 593 +/-23 nm and in the FAM channel with λ_exc_ = 469 +/-20 nm and λ_em_ = 525.5 +/-23.5 nm. Image acquisition was controlled with the Metamorph software (5 min/frame, 100 ms exposure time) for 500 min or 90 min for the controls and analysed using the Fiji software. The integrated fluorescence intensities divided by the volume of the droplet as a function of time was obtained for at least twelve coacervates microdroplets

## Author Contributions

HM, MK, TYDT conceived the project; TYDT, HM, BD, JMIA, KLV, VM undertook the experiments and data analysis; HM, TYDT, BD, JMIA, KLV, KJ discussed the results; TYDT, HM, BD, KLV wrote the manuscript.

## Notes

The authors declare no competing financial interests.

## Acknowledgment

Funding was provided by the Volkswagen Foundation and the MaxSynBio consortium which is jointly funded by the Federal Ministry of Education and Research of Germany and the Max Planck Society. We thank the light microscopy facility (LMF) for their continuous support and guidance during the FRAP experiments. We acknowledge the Boehringer Ingelheim Fonds for the award of a PhD fellowship to JMIA. We thank Andre Nadler and Carl Modes for useful discussion.

